# Assembly and Interfacial Mechanics of Recombinant Surfactant Protein B in Mixed Detergent Systems and Lipid Monolayers

**DOI:** 10.64898/2026.09.04.743498

**Authors:** Tadiwos Getachew Asrat

## Abstract

Although surfactant protein-B (SP-B) is essential for pulmonary surfactant function, the experimental structure of mature SP-B remains undetermined, and the molecular basis of its function is unclear. Here, we characterize the structural and functional behavior of a recombinant SP-B construct (rSP-B) in DPC/SDS detergent micelles and DPPC/POPG lipid monolayers. Light scattering showed that dilute DPC/SDS (9:1, 0.2% w/v) is intrinsically heterogeneous, containing both micellar (~4 nm) and supramicellar structures (~68–400 nm), consistent with non-ideal surfactant mixing at low detergent concentration. Dialysis resulted in a dominant, reproducible mesoscale structure (~600 nm) in which rSP-B retained predominantly *α*-helical structure (~47%), whereas 200 nm filtration removed most of the protein and helical signal, suggesting that rSP-B is associated with the large assemblies, although the resulting complex sizes exceed the regime typically accessible to solution NMR. In separate detergent-only measurements, higher detergent concentrations (0.5–2% w/v) gave monomodal micelles (3.6–4.1 nm), suggesting that detergent concentration influences assembly size. In parallel, incorporation of rSP-B into DPPC/POPG monolayers stabilized the hysteretic response and attenuated cycling-induced area loss relative to protein-free films, with compressibility signatures consistent with out-of-plane deformation and reversible reorganization of collapsed material. These results inform strategies for optimizing detergent conditions for future structural studies and support a model in which rSP-B may promote retention and reversible reorganization of collapsed interfacial material, potentially through mechanically coupled reservoirs.

## 1. Introduction

Pulmonary surfactant (PS) is a lipoprotein complex that lines the alveoli and lowers surface tension at the air–liquid interface, preventing alveolar collapse during expiration [1]. Impaired surfactant function contributes directly to severe pulmonary disorders, including neonatal respiratory distress syndrome (NRDS) and acute respiratory distress syndrome (ARDS) [2, 3]. Although exogenous surfactant therapy is highly effective in premature infants with NRDS, its limited success in adult ARDS highlights the need for improved therapeutic strategies [4]. However, the molecular mechanisms by which surfactant proteins organize lipid membranes to achieve and maintain ultra-low surface tension during dynamic respiratory cycles remain incompletely understood [5]. Defining the structural organization of PS is therefore critical for understanding surfactant dysfunction in lung injury and for guiding the development of more effective therapies.

The unique biophysical properties of PS arise from its highly specialized molecular composition, consisting primarily of phospholipids (~90% by mass) and a smaller fraction of surfactant-associated proteins (SPs) [6]. The dominant lipid species, DPPC, forms the condensed phase responsible for near-zero surface tension during compression. The hydrophilic proteins SP-A and SP-D primarily contribute to innate host defense, whereas the hydrophobic proteins SP-B and SP-C facilitate lipid adsorption, film spreading, and interfacial stabilization [7]. Among these components, SP-B is uniquely indispensable. Beyond lipid reorganization, SP-B contributes to lamellar body formation, surfactant secretion, SP-C processing, and tubular myelin assembly [8–11]. Consistent with these roles, congenital SP-B deficiency is lethal, whereas SP-C deficiency allows postnatal survival with progressive lung disease [12, 13]. Collectively, these observations identify SP-B as a central determinant of surfactant biogenesis and function.

SP-B is synthesized in alveolar type II cells as a large precursor that undergoes proteolytic processing to yield a mature 79-residue (~8.7 kDa) peptide [14]. The mature protein is highly hydrophobic and cationic, properties that promote membrane insertion and interactions with anionic phospholipids. SP-B belongs to the saposin-like protein (SAPLIP) family, whose members share a conserved helical fold stabilized by three intrachain disulfide bonds [15]. A distinctive feature of SP-B is an additional cysteine that forms an interchain disulfide bond, producing a covalent homodimer unique within the SAPLIP family [16]. Although monomeric SP-B retains partial activity, the dimeric form is required for optimal pulmonary function *in vivo* [17, 18]. Homology models predict four to five amphipathic helices per monomer, including a proline-rich N-terminal segment implicated in membrane insertion and curvature modulation [19, 20]. Despite its central biological role, no high-resolution experimental structure of full-length SP-B currently exists. NMR studies of truncated constructs such as Mini-B resolve elements of the saposin fold in detergent micelles; however, these simplified models do not fully reproduce the oligomeric organization, membrane topology, and dynamic lipid interactions of full-length SP-B [15, 21]. Consequently, the structural mechanisms by which SP-B organizes surfactant membranes remain poorly defined.

Determining SP-B structure has been challenging due to its intrinsic biophysical properties. Its amphipathic, membrane-disruptive nature complicates heterologous expression and *in vitro* refolding, as correct folding requires membrane insertion and proper disulfide pairing [22–25]. Structural studies therefore require membrane-mimetic environments that stabilize SP-B while remaining compatible with high-resolution spectroscopy. Solution NMR, in particular, requires small, monodisperse protein–detergent complexes (PDCs) that undergo rapid isotropic tumbling [26]. While lipid vesicles better approximate native membranes, their size restricts NMR analysis primarily to solid-state approaches [27]. In contrast, detergent micelles and isotropic bicelles support high-resolution NMR measurements provided the resulting PDCs remain small and homogeneous, minimizing anisotropic interactions [28, 29]. Achieving these conditions can be challenging for SP-B. Its strong membrane affinity and tendency to self-associate can produce large or heterogeneous PDCs that compromise spectral resolution. Reliable structural characterization therefore requires careful control of PDC size, composition, and protein integrity.

While detergent micelles provide a tractable environment for structural characterization, they do not reproduce the interfacial environment in which pulmonary surfactant functions. Lipid monolayers at the air–water interface therefore serve as a complementary biophysical model for evaluating surfactant protein activity under controlled compression–expansion cycles that mimic respiratory mechanics [6, 7]. Interfacial properties can be quantified *in vitro* using Langmuir trough measurements, including surface pressure–area isotherms, hysteresis, and collapse–recovery dynamics in defined lipid systems [30–32]. Consequently, monolayer studies provide a sensitive platform for relating protein-induced changes in film behavior to surfactant function and for evaluating whether recombinant SP-B retains interfacial activity.

In previous work, we established a method for recombinant production, purification, and *in vitro* folding of SP-B [33, 34]. To reduce host toxicity and aggregation during expression, a modified construct lacking the insertion sequence and Cys48 (hereafter rSP-B) was used. Among the membrane-mimetic systems examined, mixed DPC/SDS micelles restored substantial α-helical structure comparable to that reported for native SP-B isolated from lung lavage or animal-derived surfactant preparations (Figure 3B) [35, 36], supporting its utility as a practical system for probing the biophysical principles that govern SP-B–membrane interactions. Here, we investigate rSP-B in DPC/SDS micelles and DPPC/POPG lipid monolayers to address two open questions: first, whether membrane-mimetic conditions can be tuned to yield PDCs suitable for high-resolution structural analysis, and second, whether rSP-B retains surfactant-relevant interfacial activity outside the detergent environment. Together, these complementary systems show that rSP-B retains substantial α-helical secondary structure in dilute DPC/SDS before and after dialysis and exhibits surfactant-relevant interfacial activity in detergent-free lipid monolayers. These findings provide a basis for further optimization of detergent conditions for future structural studies and offer insight into the potential contribution of rSP-B to surfactant-film stability during cyclic compression and expansion.

## 2. Materials and Methods

### 2.1. Materials

All lipids were purchased from Avanti Polar Lipids (Alabaster, AL, USA). Detergent stocks were prepared by mixing DPC/SDS (9:1 molar ratio) in 20 mM Tris-HCl (pH 7.5) without added NaCl to a final detergent concentration of 0.2% (w/v) (~ 5.8 mM total detergent). rSP-B samples reached a final pH of ~5 through pH-gradient elution as described below. Detergent-only controls were prepared and measured at both pH 7.5 and pH 5 to assess the effect of pH on detergent self-assembly. Protein concentration was determined using the Bradford assay. DPPC/POPG (7:3, w/w) lipid mixtures were dissolved in chloroform/methanol to a final concentration of 1 mg/mL.

### 2.2. Protein Expression and Purification

rSP-B was expressed from pET11a-SN-SP-B in *E. coli* C43 (>1 ×10^6^ cfu/μg DNA) grown in 2× YT medium as described in [33]. Protein expression was induced at OD_600_ = 0.6 using 0.4 mM IPTG and continued for 3 h at 30 °C. Purification was performed using Ni^2+^ immobilized metal affinity chromatography (IMAC) on Sepharose 6 Fast Flow (GE Healthcare) in a PD-10 gravity column. Protein refolding was achieved on-column through stepwise dilution of urea from 6 M to 0 M. The protein was subsequently exchanged into the desired detergent/buffer system and eluted using a linear pH gradient (pH 7.4 to 5.0). Purified protein was used immediately or stored at 4 °C for short-term experiments.

### 2.3. Dialysis and Filtration of Protein–Detergent Complexes

Protein–detergent samples were dialyzed using regenerated cellulose membranes with a molecular weight cut-off (MWCO) of 1 kDa (Spectrum Laboratories, USA). Samples were dialyzed against 3 L of distilled water at 4 °C with continuous stirring overnight (≈ 8–12 h). Following dialysis, samples were collected and immediately analyzed.

For filtration experiments, dialysed samples were passed through a 200 nm syringe filter (Sartorius, Göttingen, Germany), and the filtrate was analyzed by DLS and CD.

### 2.4. Circular Dichroism (CD)

Secondary structure of rSP-B in 9:1 DPC/SDS micelles (0.2% w/v) was measured using a Jasco J-810 spectropolarimeter (Jasco Inc., Tokyo, Japan). Spectra were collected from 190–260 nm at a scan rate and pitch of 1 nm/s and 1 nm, respectively, using a 0.5 mm pathlength quartz cuvette at 25 °C. HT voltage remained below 600 V across the measured range. Twenty scans were averaged for each sample, baseline-corrected against buffer/detergent alone, and converted to mean residue ellipticity (MRE). Percent *α*-helicity was estimated from the normalized spectra using established empirical relationships [37].

### 2.5. Dynamic Light Scattering (DLS)

Particle size distributions of detergent solutions and rSP-B/detergent samples were measured using a Zetasizer Nano ZS (Malvern Instruments Ltd, Malvern, Worcestershire, UK) at a scattering angle of 173° and 25 °C, with a 2 min temperature equilibration prior to each measurement. Dispersants and solvents were filtered through sintered glass filters (Fisherbrand); protein-containing samples were not filtered prior to measurement, except where filtration was the variable under study. The viscosity and refractive index of the dispersant medium were used for size calculation. Each sample was measured three times, and results were confirmed across six independent preparations. The hydrodynamic diameter (*d*_*h*_) was calculated from the translational diffusion coefficient (*D*_*t*_) using the Stokes–Einstein equation (Eq. 1):

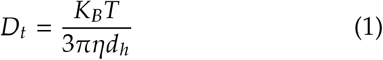

where *K*_*B*_ is Boltzmann’s constant, *T* is the absolute temperature, and *η* is the viscosity of the medium.

Data were analyzed using Malvern Zetasizer software (v7.03). Particle sizes were additionally validated by fitting the raw autocorrelation function (ACF) to a sum of exponential decays (IGOR Pro 7) and comparing the results with the Zetasizer NNLS-derived distribution.

### 2.6. Nanoparticle Tracking Analysis (NTA)

Particle size distributions of larger assemblies (>30 nm) were measured using a NanoSight NS500 (Malvern Instruments Ltd, Malvern, Worcestershire, UK) equipped with a 532 nm laser at room temperature (25 °C). Samples were analyzed undiluted. Videos of 90 s were recorded at 30 frames per second (camera shutter 2.48 ms, camera gain 200) and analyzed using NTA software (version 2.3, build 0034) with a detection threshold of 10 and a viscosity of 0.89 cP. The translational diffusion coefficient was determined from the two-dimensional mean square displacement (⟨*x, y*⟩^2^) according to Eq. 2:

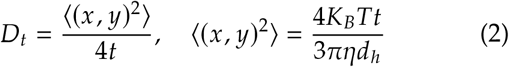

where *t* = 1 / FPS and other variables are as defined above.

### 2.7. Langmuir Trough Measurements

Surface pressure–area (Π–*A*) isotherms were recorded using a Kibron MicroTrough XS (59 × 208 mm; Kibron Inc., Helsinki, Finland, software version 3.6.1) at ambient temperature (25 °C). Lipids were mixed by weight to a final concentration of 1 mg/mL in a chloroform/methanol solution. Aliquots of rSP-B at 2, 4, 6, 8, and 10 wt% (with respect to lipid) in a 1:1 (v/v) chloroform/methanol mixture were added to the 1 mg/mL lipid solution. Ten μL of lipid–protein solution was spread onto the air–water interface of ultrapure water (18.2 MΩ·cm) using a microsyringe and allowed to equilibrate for at least 10 min prior to compression. For cyclic isotherms, a single continuous measurement captured both compression and expansion phases across all cycles, performed at ambient temperature. Local slopes of the expansion branch were obtained by linear regression of Π against *A* within 5 mN/m pressure windows.

Two-dimensional isothermal compressibility modulus 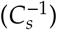 was calculated as shown in Eq. 3:

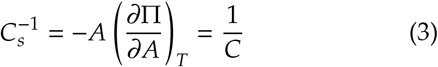

where *A* is the monolayer area, Π is the surface pressure, and *C* is the monolayer compressibility. Data were smoothed and analyzed using IGOR Pro 7 (Wavemetrics, Portland, OR, USA).

The hysteresis area (*H*) was calculated according to Eq. 4:

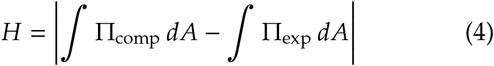

where the compression and expansion branches were interpolated onto a common area grid spanning the range over which both branches had data, and the integrals were evaluated using the trapezoidal rule. Normalized hysteresis was calculated as *H* / *W*_comp_ ×100, where *W*_comp_ is the compression-branch work evaluated over this same shared area range.

## 3. Results

### 3.1. Dilute DPC/SDS is heterogeneous in the absence of rSP-B

Dynamic light scattering (DLS) was used to analyze the size distribution of rSP-B in 0.2% (w/v) DPC/SDS (9:1) micelles prior to dialysis (Figure 1). The particle size distribution (PSD) was highly polydisperse and multimodal, with three principal populations observed across independent preparations. A dominant peak at ~ 4 nm was consistent with spherical DPC/SDS micelles. Two broader, partially overlapping populations spanning ~ 68– 400 nm were also detected, indicating the presence of larger assemblies. A minor apparent population at ~ 5– 6 μm was also reported by the software; however, analysis of the raw ACF indicated that this feature arose from the noise-dominated long-lag region and was not reliably resolved. Multi-exponential fitting of the autocorrelation function confirmed particle sizes in good agreement with the NNLS-derived distribution (Table S1). A similar PSD was observed in detergent-only controls (Figure S1), including the presence of larger size populations, suggesting that this heterogeneity reflects intrinsic surfactant self-assembly rather than protein-induced aggregation. Under these conditions, association of rSP-B with specific particle populations was not resolved.

**Figure 1.**
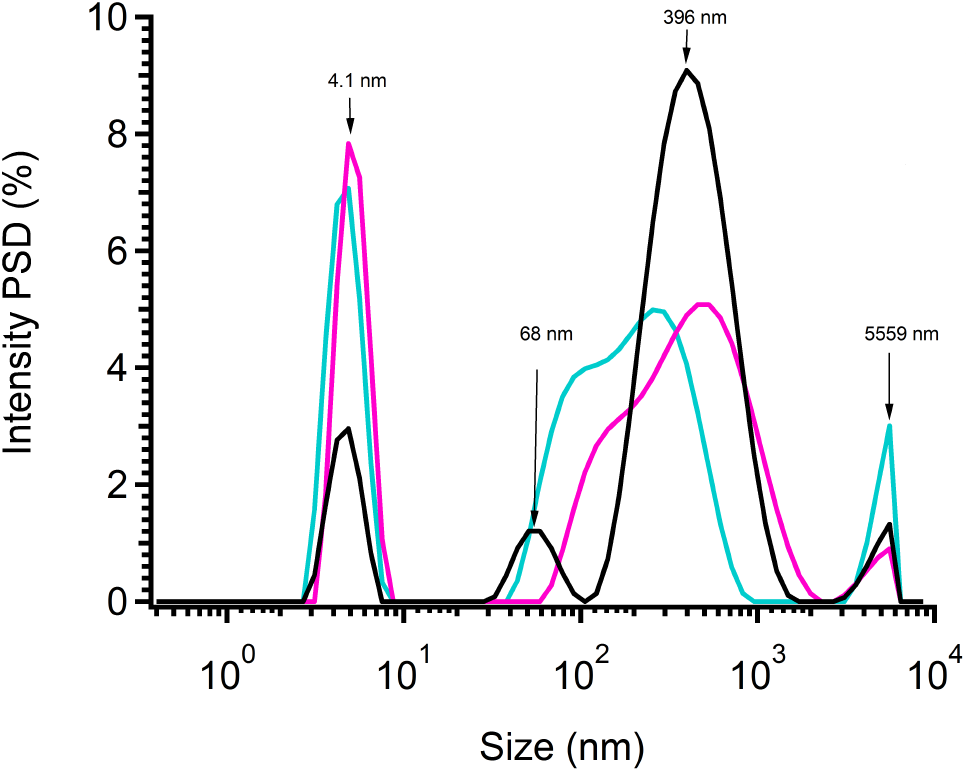
Particle size distribution of rSP-B in 0.2% (w/v) DPC/SDS (9:1) micelles measured by dynamic light scattering prior to dialysis, showing a micelle-sized population at ~4 nm, broader particle populations at ~68–400 nm, and a minor population at ~5–6 μm.

### 3.2. Supramicellar assemblies detected by NTA

Nanoparticle tracking analysis (NTA) was used to characterize larger particles in the rSP-B/DPC/SDS system (Figure 2). Due to its limited detection range for small particles (~ 30 nm) [38], NTA did not detect the ~4 nm micellar population observed by DLS. Instead, a consistent population with a modal diameter of ~ 120 nm was observed across independent measurements. Approximately 90% of particles were below ~ 450 nm, with a distribution spanning ~ 60–500 nm. These measurements overlaps the larger hydrodynamic-size populations detected by DLS, supporting the presence of mesoscale particles in this size range.

**Figure 2.**
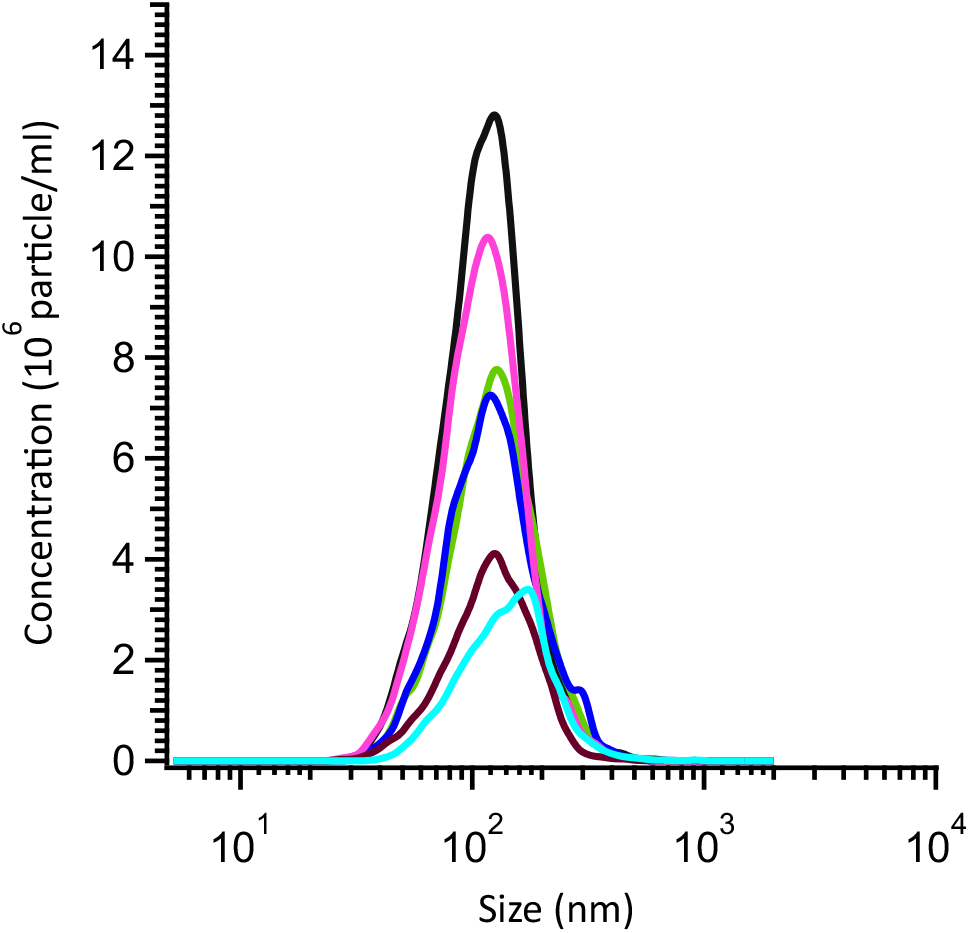
Particle size distribution of rSP-B in 0.2% (w/v) DPC/SDS (9:1) measured by nanoparticle tracking analysis showing particles in the ~60–500 nm range with a modal diameter of ~120 nm (n = 6 independent measurements). A representative NTA video illustrating the Brownian motion of the tracked particles is provided as supplementary Video S1.

### 3.3. Dialysis yields a single dominant population of large assemblies

Dialysis of the rSP-B/DPC/SDS system using a 1 kDa molecular weight cutoff membrane resulted in a marked redistribution of particle sizes (Figure 3A). Following dialysis, the ~ 4 nm population was no longer detected, and the PSD shifted to a single, unimodal peak centered at ~ 600 nm. This distribution was reproducible across independent preparations and consistent across intensity-, volume-, and number-weighted analyses. ACF fitting also indicated a predominantly unimodal population, and the cumulant PDI was ≈ 0.45 (Figure S5). The dialyzed sample retained predominantly *α*-helical signal (Figure 3B). The translational diffusion coefficient from the manual ACF fit decreased from ~ 1.37 ×10^−^12 to 9.75 ×10^−^13 m^2^ s^−1^ (modal diameters ~320 and ~ 450 nm by this fit, and ~ 400 and ~ 600 nm by NNLS; Table S1). The loss of the micellar population, despite its apparent size exceeding the membrane cutoff, suggests that micelle stability is disrupted during dialysis. Filtration of the dialyzed rSP-B/DPC/SDS sample through a 200 nm membrane reduced the derived count rate, shifted the diffusion coefficient back toward the pre-dialysis scale, and caused a substantial loss of *α*-helical CD signal, whereas rSP-B in methanol was unaffected. The larger populations in the detergent-only blank persisted after the same filtration. Together, these observations suggest that the helical signal depends on intact large assemblies, although direct association of rSP-B with these particles was not resolved.

**Figure 3.**
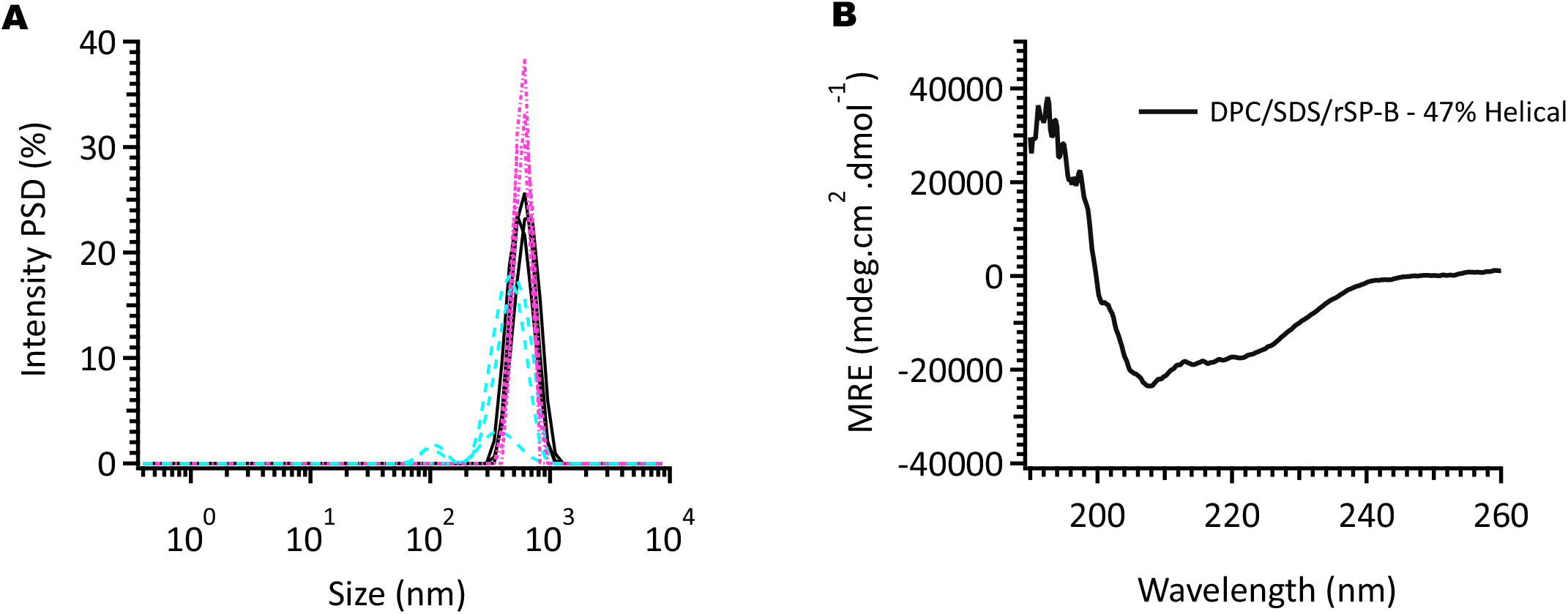
Effect of dialysis on rSP-B in 0.2% (w/v) DPC/SDS (9:1). (A) Particle size distribution measured by dynamic light scattering following dialysis, showing loss of the ~4 nm population and emergence of a single distribution centered at ~600 nm. (B) Circular dichroism spectra of dialyzed rSP-B/DPC/SDS samples indicating predominantly *α*-helical secondary structure.

### 3.4. Stabilized hysteretic response and collapse behavior in rSP-B–lipid monolayers

To assess whether rSP-B retains functional activity independent of the detergent environment, its interfacial behavior was evaluated in a detergent-free lipid monolayer system. Cyclic Π–A compression–expansion isotherms of the DPPC/POPG (7:3) film containing 6 wt% recombinant SP-B showed isotherms retracing a similar hysteresis loop across cycles 4–6 (Figure 4), whereas the protein-free film changed progressively with cycling (Figure S2). Quantitative analysis supported this behavior: normalized hysteresis, calculated as hysteresis area normalized to compression work, remained stable across cycles 4–6 (61.8%, 61.2%, and 60.3%; range 1.5 percentage points) (Table 1). Molecular area at 10 mN/m was unchanged between cycles 4 and 6 (84.1 Å^2^), whereas areas at 30 and 50 mN/m decreased modestly by 5.7% and 7.2%, respectively, indicating some residual loss or reorganization of interfacial material during cycling. In contrast, the lipid-only control showed an increase in normalized hysteresis from 37.2% to 45.2% between cycles 4 and 8 despite little change in compression work, accompanied by a ~25% contraction in molecular area at 10, 30, and 50 mN/m (Table 1), indicative of progressive film reorganization. Notably, normalized hysteresis was higher in the rSP-B film than in the lipid-only film. Thus, rSP-B did not reduce hysteresis magnitude but stabilized the hysteretic response and attenuated, without eliminating, cycling-induced area loss. A distinct change in slope was observed in the expansion branch of the rSP-B film near Π ≈ 25 mN/m: the local slope fell 4–7-fold between the 30–25 and 25–20 mN/m windows in each of cycles 4–6 (e.g., ^−^2.2 to ^−^0.35 mN/m per Å^2^/molecule in cycle 4), whereas the protein-free film showed no comparable break (cycle 4: ^−^0.67 to ^−^0.60; cycle 8: ^−^0.55 to ^−^0.80 (mN/m)/(Å^2^/molecule)). This feature is compatible with reversible folding collapse and reincorporation of material excluded at the plateau onset during compression (Π ≈ 62 mN/m). In line with previous reports [39], respreading was facilitated in the liquid-expanded (LE) phase, where available interfacial area may promote reincorporation of collapsed material.

**Table 1.**
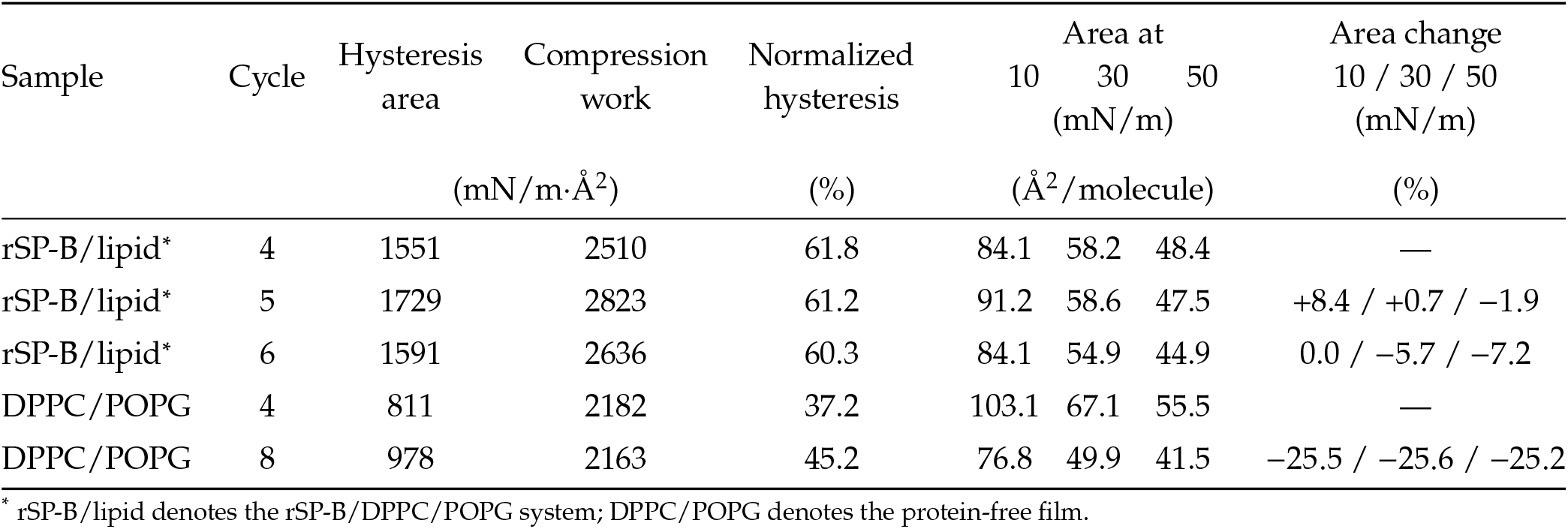
Cycle analysis of DPPC/POPG (7:3) monolayers with 6 wt% rSP-B and of protein-free DPPC/POPG films. Hysteresis area and compression work are reported for the indicated cycles, together with normalized hysteresis. Molecular area is given at 10, 30, and 50 mN/m on the compression branch. Area change (%) is calculated relative to cycle 4 at the corresponding surface pressure. The last cycle analyzed is cycle 6 for the rSP-B film and cycle 8 for the protein-free film.

**Figure 4.**
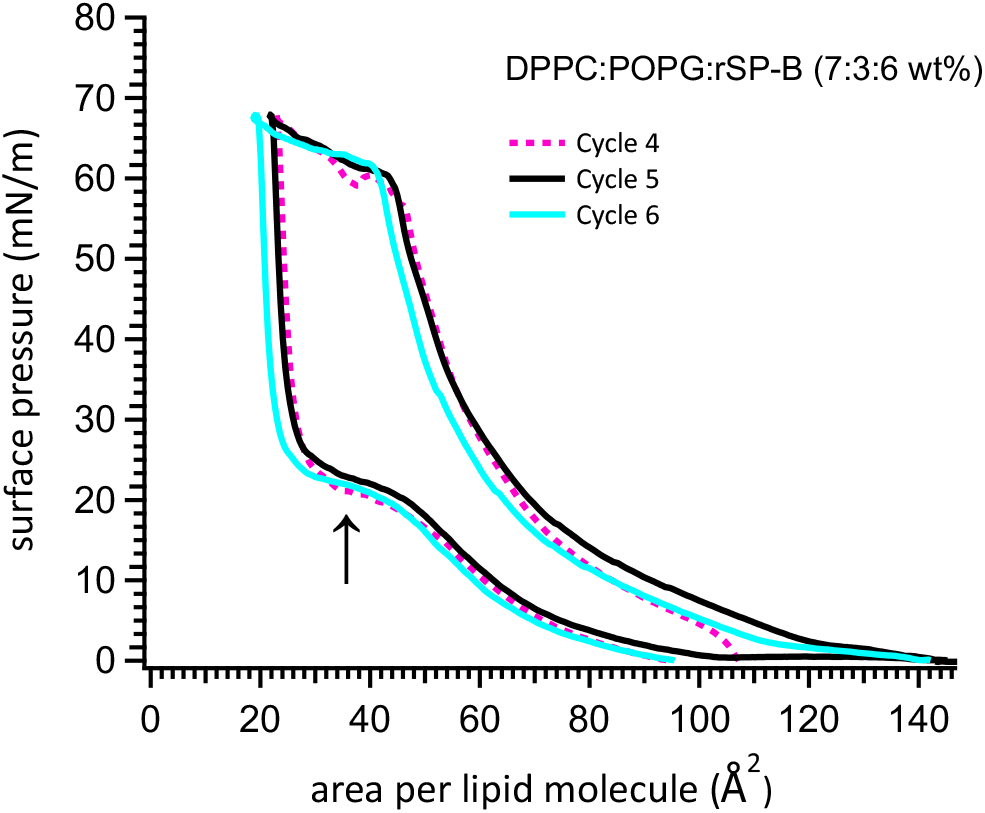
Cyclic surface pressure–area (Π–A) isotherms of a DPPC/POPG (7:3) monolayer containing 6 wt% rSP-B measured under repeated compression–expansion cycles. Successive cycles (4–6) show closely similar hysteresis loops. A change in slope is observed in the expansion branch at Π ≈ 25 mN/m, and the compression branch reaches a plateau near Π ≈ 62–68 mN/m.

Analysis of the compressibility modulus 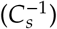 as a function of mean molecular area (MMA), derived from the Π–A isotherms (Figure 5), shows that at MMA ≈ 50 Å^2^ the monolayer is in the liquid-condensed (LC) phase 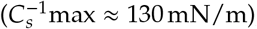, while compressibility (*C*) sharply increases through the transition region (corresponding to Π ≈ 56–61 mN/m, discussed below), then peaks near MMA ≈ 40 Å^2^ and remains nearly constant as surface pressure rises slightly and approximately linearly from the collapse onset (Π ≈ 62 mN/m) to Π 68 mN/m, at which point the compression cycle was reversed. Upon expansion, the film transiently enters a highly ordered, solid-like state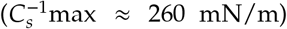, followed by a rapid transition into the liquid-expanded (LE) phase 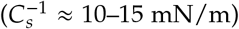 with minimal change in molecular area (≈ 5 Å^2^), accompanied by a sharp drop in surface pressure from ≈ 68 to 25 mN/m (Figure 4). The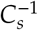 –Π relationship (Figure S3) shows maximal lipid packing at Π ≈ 55 mN/m, consistent with the MMA ≈ 50 Å^2^ packing maximum described above. Between Π ≈ 56–61 mN/m, an increase in compressibility despite rising surface pressure suggests partial relaxation of lateral stress, potentially via out-of-plane deformation such as buckling.

**Figure 5.**
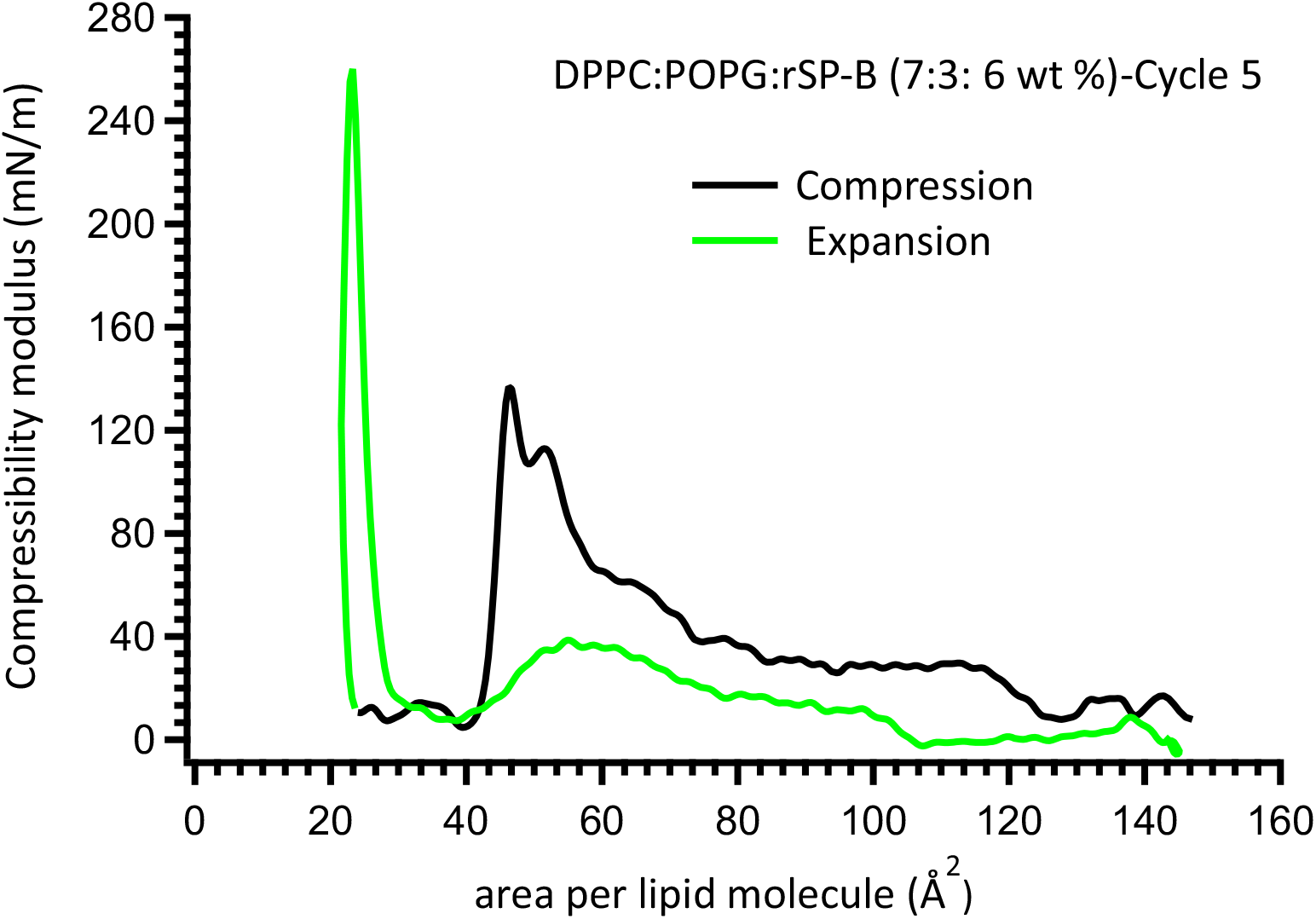
Compressibility modulus 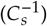 of a DPPC/POPG (7:3) monolayer containing 6 wt% rSP-B derived from surface pressure–area (Π–A) isotherms as a function of mean molecular area (MMA). The system exhibits 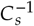 values ranging from approximately 10 to 260 mN/m across compression and expansion cycles.

## 4. Discussion

### 4.1. PDC assembly and implications for NMR sample design

DPC/SDS mixtures are widely used as membrane mimetics [40]. Under dilute conditions (0.2% (w/v) DPC/SDS, 9:1, Tris-HCl), however, the system does not adopt a uniform micellar state. Instead, DLS reveals a heterogeneous, multimodal particle distribution containing both ~ 4 nm particles and larger mesoscale populations (Figure 1). The ~ 4 nm species agrees with previous reports for DPC/SDS mixed micelles [41], whereas the persistence of larger species in detergent-only controls (pH 7.5) (Figure S1) and after filtration indicates that the observed heterogeneity arises from intrinsic surfactant self-assembly rather than protein-induced aggregation or sample handling artifacts. Mixed ionic–zwitterionic surfactant systems commonly show non-ideal mixing, in which micelle composition and aggregation behavior depend on total surfactant concentration, electrostatic interactions, and ionic screening [40, 42]. Such effects may contribute to the heterogeneity observed here at low detergent concentration, although other contributors, including trace large particles, cannot be excluded. Within this regime, rSP-B likely partitions among coexisting assemblies, although preferential localization cannot be resolved from the present data.

Dialysis substantially reorganized this assembly landscape. Following dialysis, the ~4 nm population decreases and the system reorganizes into a reproducible mesoscale structure centered at ~600 nm (Figure 3A). The emergence of a defined population argues against stochastic aggregation and instead suggests selection of a preferred structure under monomer-limited conditions. Given rapid monomer–micelle exchange (*τ*_1_) [43] relative to slower micelle restructuring and relaxation (*τ*_2_) [44], monomer depletion during dialysis is expected to suppress high-curvature micelles and bias the system toward lower-curvature or kinetically trapped species over the experimental timescale (~ 8 h). The resulting structures may reflect either a lower-curvature free-energy minimum or kinetic stabilization arising from arrested coarsening under conditions of limited monomer exchange and energetic barriers to curvature reorganization. Circular dichroism shows that rSP-B retains native-like secondary structure within these assemblies (Figure 3B). Conversely, filtration (200 nm)-induced disruption of the mesoscale structures in the dialysed sample compromised protein secondary structure. rSP-B in methanol remained structurally intact under comparable conditions, suggesting that these structures contribute to structural stabilization. The retention of native-like secondary structure supports a role for rSP-B in interfacial packing, although passive partitioning into pre-existing structures without substantial perturbation of assembly energetics cannot be excluded. Regardless of origin, the observed mesoscale assembly, though comprising a single dominant population, remains substantially larger than the small complexes exhibiting rapid isotropic tumbling generally required for solution NMR.

DPC/SDS self-assembly appears to be concentration dependent. In separate detergent-only measurements in the same buffer, DLS of DPC/SDS at 0.5–2% (w/v) showed a single monomodal population with modal hydrodynamic diameters of 3.6–4.1 nm (n = 3 per concentration; Figure S4), consistent with small mixed micelles and in contrast to the multimodal distribution of the 0.2% (w/v) detergent-only blank. At these concentrations, the system likely resides further above the effective CMC, where increased surfactant availability and rapid monomer–micelle exchange facilitate micellar equilibration and favor more uniform micelles [45, 46]. In contrast, the more dilute 0.2% (w/v) solution yielded the heterogeneous, multimodal populations described above (Figure 1). These observations suggest that DPC/SDS assembly in this buffer can be tuned through changes in surfactant concentration, giving access to distinct organizational states in detergent-only samples. This concentrationdependent behavior provides a starting point for optimizing DPC/SDS conditions for future rSP-B studies and highlights how variations in detergent formulation can influence mesoscale organization in mixed surfactant systems used for membrane-protein biophysics.

The concentration-dependent organizational behavior observed here has implications for solution NMR of membrane proteins. In dilute DPC/SDS systems, ionic strength is expected to modulate micelle–aggregate equilibria through electrostatic screening of SDS headgroups, thereby influencing surfactant self-assembly behavior [40, 47]. However, increased ionic strength can compromise NMR sensitivity through increased sample conductivity, associated RF losses, and enhanced sample heating [48, 49]. The system therefore presents a potential trade-off between micellar homogeneity and NMR performance. In the 20 mM Tris-HCl buffer used here (ionic strength ~16 mM, no added NaCl), increasing detergent concentration alone produced monomodal micelles in detergent-only samples, although whether rSP-B forms small, monodisperse complexes under these conditions remains to be tested. Added salt may also shift micelle– aggregate equilibria, but its effect on this system was not tested and could favor either smaller or larger assemblies; any such benefit would have to be weighed against the NMR costs noted above. Identifying conditions that balance size homogeneity, protein stability, and NMR sensitivity therefore requires systematic optimization of detergent concentration and buffer composition.

### 4.2. Mechanistic basis of rSP-B mediated collapse reversibility

Efficient recovery of surfactant material following collapse is essential for maintaining low surface tension during respiratory cycling, and our results support a contribution of rSP-B to this process. Highly condensed monolayers typically collapse through fracture, whereas fluid lipid films collapse through solubilization or material exclusion, both of which can result in irreversible loss of interfacial material [30]. In contrast, PS exhibits a laterally heterogeneous architecture in which condensed domains are embedded within a continuous liquid-expanded (LE) matrix. This organization accommodates compressive stress through buckling and folding while preserving the interfacial continuity required for rapid respreading [50]. However, protein-free DPPC/POPG films showed a progressive increase in hysteresis and contraction of molecular area during repeated compression and expansion cycles (Figure S2, Table 1), a pattern suggestive of progressive loss of interfacial material. These changes indicate that lipid phase coexistence alone is insufficient to ensure reversible collapse. Incorporation of rSP-B (6 wt%) stabilized the hysteretic response across cycles 4–6 (Figure 4) and attenuated cycling-induced area loss relative to the protein-free film. These findings suggest that rSP-B contributes to collapse recovery through mechanisms that extend beyond lipid phase coexistence alone, possibly enabling more efficient preservation and restoration of interfacial material during cycling.

The stabilized hysteretic response likely reflects dynamic coupling of rSP-B to the evolving mechanical state of the monolayer during cyclic compression and expansion. SP-B preferentially partitions into the LE phase and co-redistributes with fluid lipids as the monolayer is compressed toward the equilibrium spreading pressure (Π_*e*_) [51]. Above Π_*e*_, the monolayer enters a supersaturated regime in which further increases in surface pressure are dominated by lateral compression. The collapse plateau near Π ≈62, mN /m and the feature near Π ≈25, mN/ m during expansion are better attributed to reversible redistribution of material than to irreversible loss from the interface, with repeated cycling yielding a reproducible yet dissipative response (Figure 4). Classical squeeze-out models describe collapse as selective exclusion of fluid lipid components and enrichment of DPPC-rich domains at the interface [52]. More recent studies, however, demonstrate that fluid phospholipid monolayers can sustain metastable surface pressures above Π_*e*_ in captive bubble systems, indicating that material exclusion is not necessarily irreversible [53]. Our observations are therefore consistent with a modified squeeze-out framework in which material excluded during compression remains associated with the interface as folded multilayer structures or surface reservoirs that remain mechanically coupled to the interfacial monolayer [50]. Given SP-B’s known preferential partitioning into the LE phase [51], rSP-B may contribute to reversible collapse by modulating interfacial mechanics near phase boundaries, matching the deformation and retention of mechanically coupled collapsed structures inferred here.

Changes in the monolayer compressibility modulus 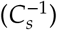, which reflects the elastic response of lipid monolayers to lateral compression, provide further insight into the mechanical basis of this recovery process [54]. Compressibility analysis indicates that maximum in-plane packing is achieved before the onset of the collapse plateau (Figure 5). Beyond this point, continued compression produces only modest increases in surface pressure. Monolayer packing reaches a maximum near Π ≈55 mN/ m, after which 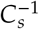 decreases (equivalently, the compressibility, 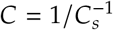, increases) despite continued elevation of surface pressure (Figure S3). These observations are consistent with the onset of out-of-plane deformation, indicative of mechanisms such as buckling and folding described in comparable systems [30], whereby further compression is accommodated through structural rearrangement rather than continued lateral condensation. Similar behavior has been reported for DPPC monolayers, where interfacial undulations emerge near Π ≈55 mN /m [55]. This mechanical behavior points to a heterogeneous interface capable of accommodating compression through curved or folded intermediates rather than catastrophic fracture.

The expansion pathway exhibits a distinct mechanical response. Although the film initially displays a rigid, transiently solid-like character, this state relaxes rapidly with minimal area expansion (Figure 5). During the initial stages of expansion, surface pressure decreases substantially despite little change in molecular area (Figure 4). This response suggests that stresses generated during collapse are relieved primarily through relaxation of collapsed structures rather than macroscopic respreading at the interface. Such behavior is indicative of a kinetically trapped state in which collapsed domains remain mechanically coupled to the monolayer and undergo structural reorganization during expansion onset. A similar persistence of metastable states during expansion has been reported for unsaturated lipid films compressed beyond Π_*e*_ in captive bubble experiments, where supercompressed monolayers do not immediately return to equilibrium until a lower surface pressure threshold is reached [56]. As expansion proceeds, the increase in 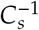 between 40 and 60 Å^2^ is consistent with progressive reinsertion and lateral redistribution of previously excluded material. In agreement, Warriner et al. [57] reported elastic redistribution of collapsed structures with limited area change before their complete reincorporation at lower surface pressures. The expansion pathway may therefore involve a two-stage recovery process: rapid stress relaxation associated with metastable collapsed structures, followed by progressive redistribution and potential reintegration of material into the interfacial film as surface pressure decreases. Collectively, these results support a model in which rSP-B may stabilize collapse intermediates that remain available for reintegration during expansion.

The results presented here support a structure–function framework for rSP-B in which detergent micellar environments influence both helical structure and the supramolecular organization of PDCs, while lipid monolayers reveal complementary interfacial activity relevant to surfactant function. In dilute DPC/SDS systems, rSP-B is present within heterogeneous detergent assemblies whose size distribution is influenced by detergent concentration and monomer availability; although protein-specific partitioning could not be resolved, rSP-B likely samples multiple membrane-associated organizational states under at this regime. Although these species are not optimal for solution NMR studies, the concentration dependence of DPC/SDS assembly behavior suggests that micellar regimes more homogeneous than those observed at 0.2% (w/v) may be accessible for rSP-B through further optimization of detergent concentration and composition. Importantly, rSP-B retains substantial *α*-helical secondary structure both before and after dialysis in dilute DPC/SDS and exhibits surfactant-relevant interfacial activity in DPPC/POPG monolayers, where its presence is associated with a reproducible cyclic response and an expansion-branch slope break compatible with reversible collapse, in contrast to the progressive changes observed in protein-free films. Together, this work provides a framework for future high-resolution structural studies aimed at defining the molecular architecture of SP-B and linking its membrane interactions to surfactant function. This is in line with the emerging view that surfactant proteins, including SP-B, may stabilize mechanically coupled interfacial reservoirs relevant to efficient surfactant recycling. In this context, recombinant SP-B provides a valuable platform for mechanistic investigations, and these findings may ultimately inform strategies for recombinant surfactant formulations.

## Supporting information

ACF_postdialysis

ACF_predialysis

## Abbreviations

PS: pulmonary surfactant
NRDS: neonatal respiratory distress syndrome
ARDS: acute respiratory distress syndrome
SP-A: surfactant protein A
SP-B: surfactant protein B
SP-C: surfactant protein C
SP-D: surfactant protein D
DPPC: dipalmitoylphosphatidylcholine
SAPLIP: saposin-like protein
rSP-B: recombinant surfactant protein B
NMR: nuclear magnetic resonance
PDC: protein–detergent complex
DPC: dodecylphosphocholine
SDS: sodium dodecyl sulfate
POPG: palmitoyl-oleoyl-phosphatidylglycerol
IMAC: immobilized metal affinity chromatography
MWCO: molecular weight cutoff
CD: circular dichroism
MRE: mean residue ellipticity
DLS: dynamic light scattering
NTA: nanoparticle tracking analysis
PSD: particle size distribution
PDI: polydispersity index
NNLS: non-negative least square
FPS: frames per second
MMA: mean molecular area
LE: liquid-expanded
LC: liquid-condensed
CMC: critical micelle concentration
RF: radio frequency.

## Acknowledgments

This study was conducted in the Department of Biochemistry at Memorial University of Newfoundland as part of my doctoral research. I thank Valerie Booth for their guidance and mentorship throughout my doctoral research and for their thoughtful review of my doctoral thesis, on which this work is based. I also thank Michael Morrow for their valuable feedback on my doctoral thesis. Financial support was provided by the Department of Biochemistry through a departmental graduate stipend, the School of Graduate Studies (SGS) fellowship, and research grant funding awarded to Valerie Booth.

## Supplementary Information

**Figure S1:**
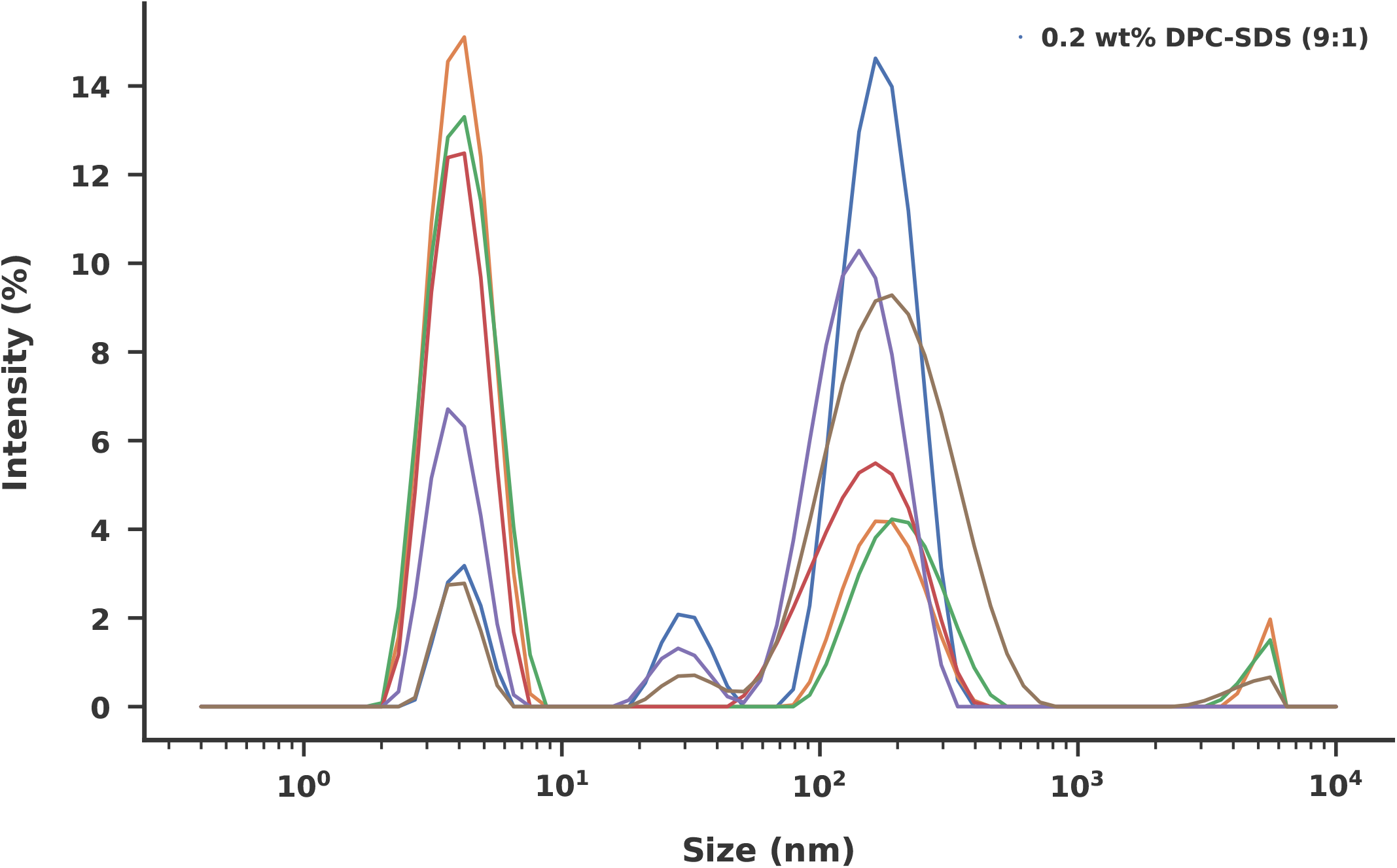
Particle size distribution of blank 0.2% (w/v) DPC/SDS (9:1) micelles measured by dynamic light scattering prior to dialysis. Six independent measurements (n = 6) show a polydisperse size distribution, reflecting the intrinsic size heterogeneity of the detergent system in the absence of rSP-B.

**Figure S2:**
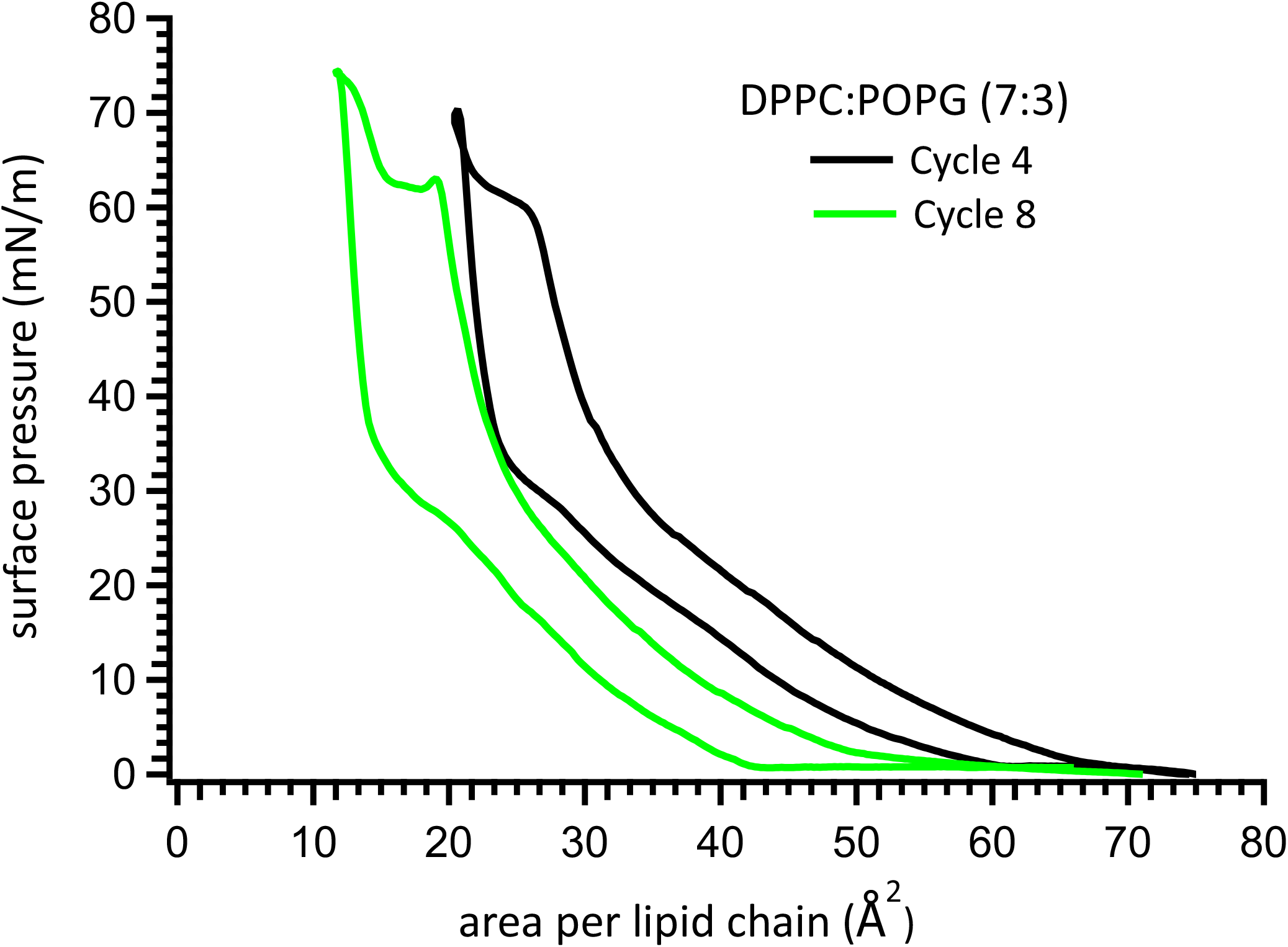
Surface pressure–area (Π–A) isotherms of the control DPPC/POPG (7:3) monolayer (without rSP-B) during repeated compression–expansion cycling. Cycles 4 and 8 are shown to illustrate the lack of a steady-state response. The pronounced shift of the cycle 8 isotherm to lower molecular areas indicates progressive loss of interfacial material upon repeated cycling.

**Figure S3:**
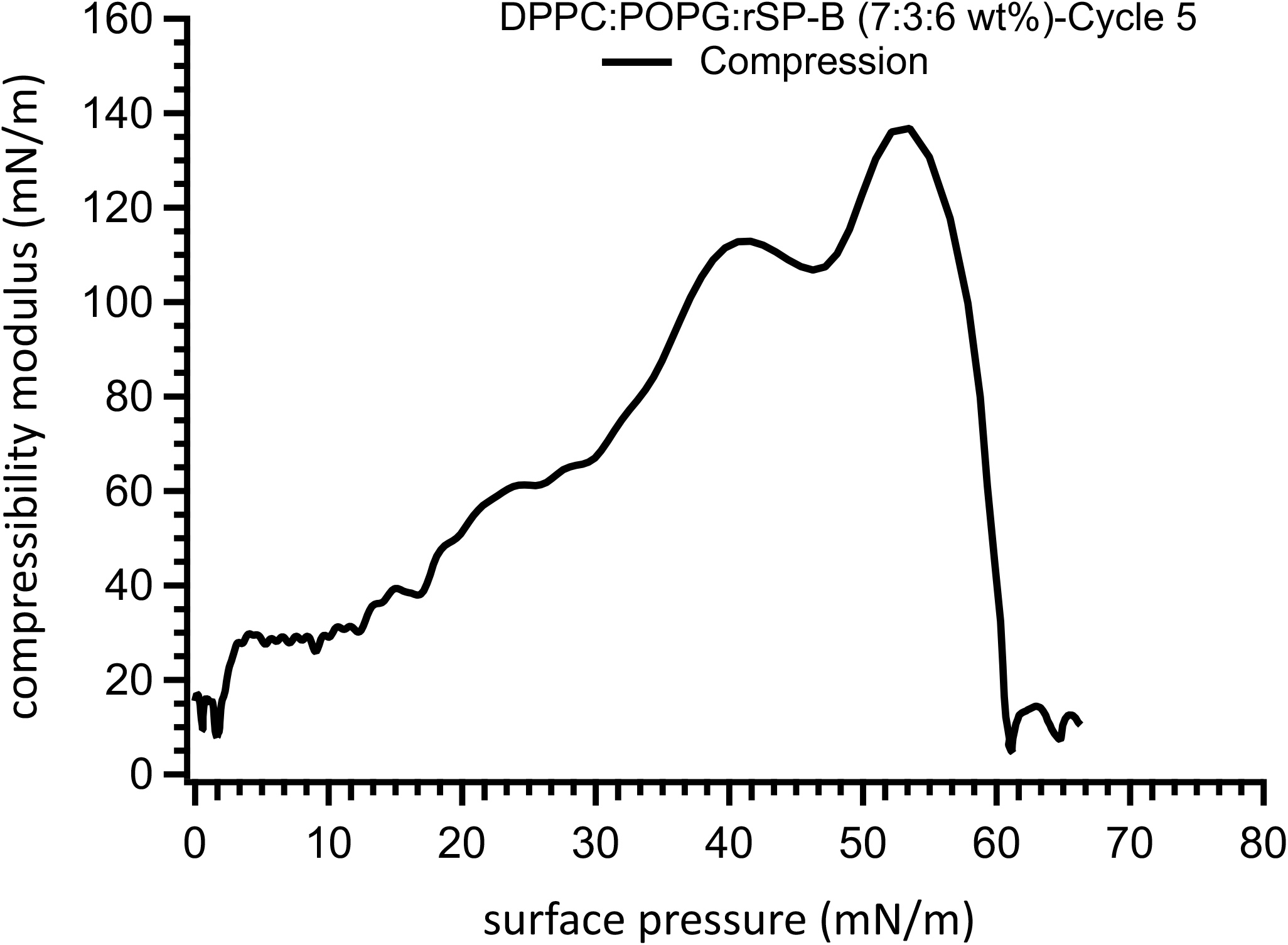
Compressibility modulus 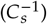 plotted as a function of surface pressure (Π) for the compression branch of a DPPC/POPG (7:3) monolayer containing 6 wt% rSP-B. At surface pressures above approximately 55 mN/m, 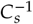 decreases with further compression, consistent with relaxation of the monolayer at high surface pressures prior to film collapse.

**Figure S4:**
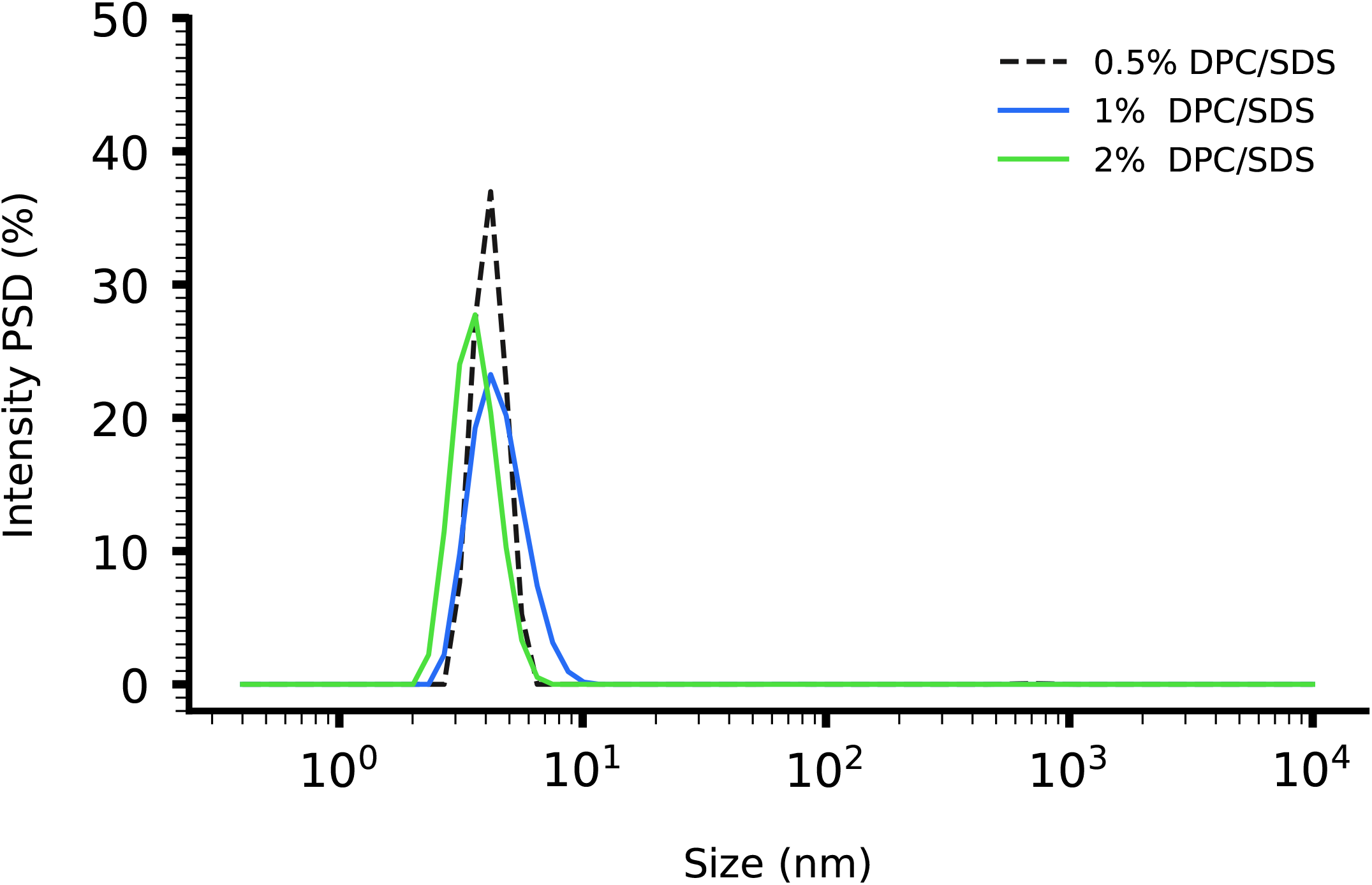
Particle size distributions of 0.5, 1, and 2% (w/v) DPC/SDS (9:1) detergent-only mixed micelles measured by dynamic light scattering. At all three concentrations, a single monomodal population is observed, with modal hydrodynamic diameters of 4.1, 4.1, and 3.6 nm, respectively (n = 3 per concentration). Intensity-, volume-, and number-weighted distributions give consistent sizes, indicating a narrow size distribution, in contrast to the multimodal distribution at 0.2% (w/v).

**Figure S5:**
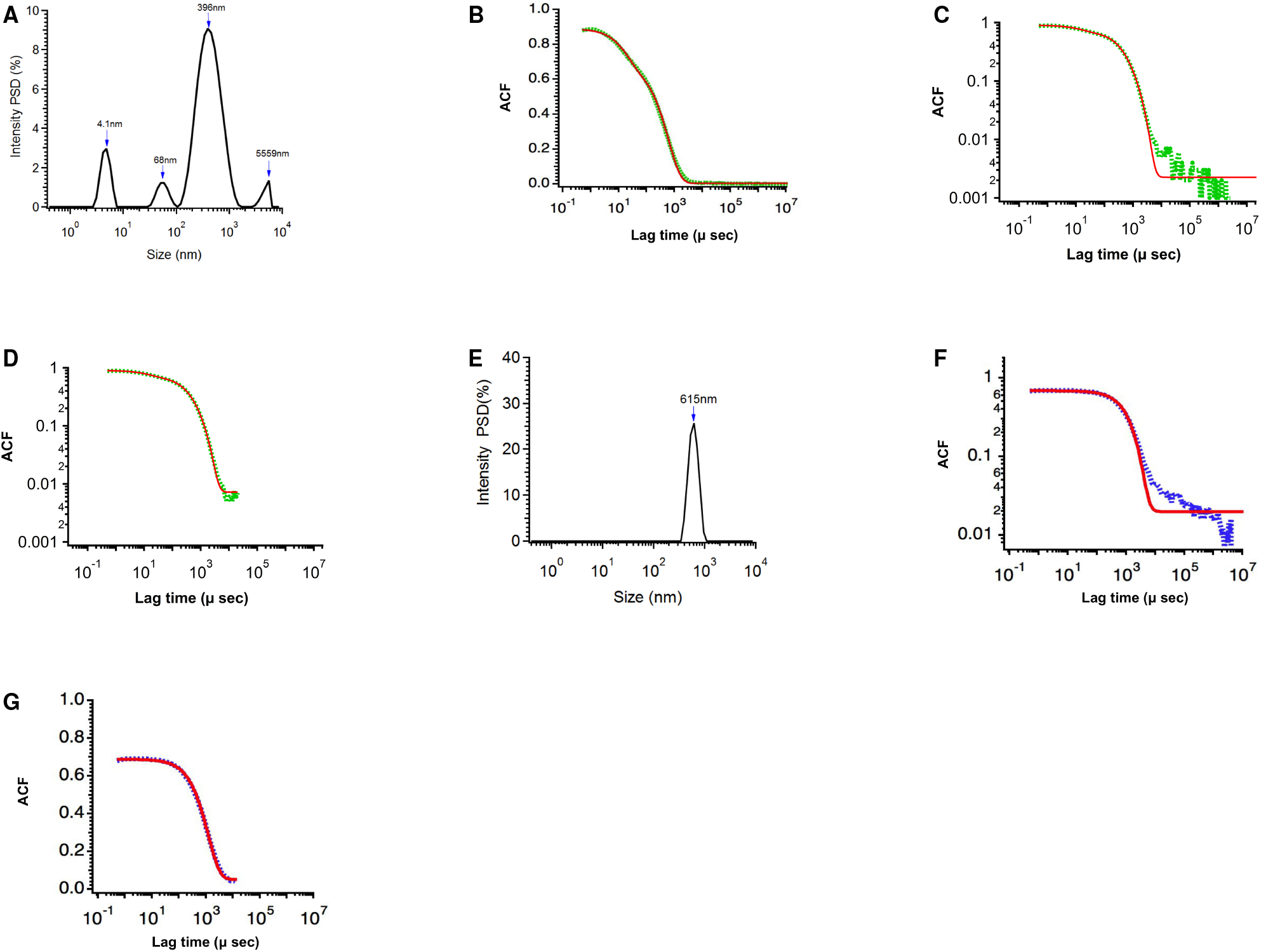
Manual exponential fits to the dynamic light scattering autocorrelation function (ACF) of rSP-B/DPC/SDS before and after dialysis. (A–D) Before dialysis. (A) Intensity-weighted particle size distribution reported by the Malvern software. (B) Measured autocorrelation function (green) and manual triple-exponential fit (red) shown on semi-logarithmic axes. The fit appears to agree well with the data. (C) The same data and fit shown on log–log axes, revealing a deviation between the fit and the data at long lag times. (D) Fit obtained after excluding the noisy long-lag-time data, where the ACF reached the noise floor; this exclusion allowed the fit to converge. The apparent particle size of ≈5560 nm reported by the Malvern software originates from this noisy region and is therefore not reliably resolved. (E–G) After dialysis. (E) Intensity-weighted particle size distribution reported by the Malvern software. (F) Autocorrelation function (blue) and single-exponential fit (red) shown on log–log axes. The data are well described by a single exponential decay. (G) Fit shown in (F) after excluding the long-lag-time noise using the same criterion, with the cutoff determined separately for this dataset at the onset of the noise. Manual fits were performed in IGOR Pro 7.

**Table S1:** Comparison of particle sizes and translational diffusion coefficients (*D*_*t*_) for rSP-B/DPC/SDS samples before and after dialysis, obtained from exponential fitting of the autocorrelation function (ACF) and from the Malvern Zetasizer NNLS algorithm.

| Sample | Method | Decay rate ($1/\mu\text{s}$) | $D_t$ ($\text{m}^2\text{s}^{-1}$) | Diameter (nm) |
| --- | --- | --- | --- | --- |
| Pre-dialysis | Full-range fit | $0.0688 \pm 0.00139$ | $8.24 \times 10^{-11}$ | 5.30 |
| | | $0.0029 \pm 0.00009$ | $3.50 \times 10^{-12}$ | 125 |
| | | $0.0009 \pm 0.00003$ | $1.13 \times 10^{-12}$ | 386 |
| | Restricted-range fit | $0.0710 \pm 0.00130$ | $8.51 \times 10^{-11}$ | 5.13 |
| | | $0.0035 \pm 0.00014$ | $4.20 \times 10^{-12}$ | 104 |
| | | $0.0011 \pm 0.00027$ | $1.37 \times 10^{-12}$ | 319 |
|  | Malvern NNLS | – | – | 4.10 (2.60–7.50); 68 (37–122); 396 (141–1281) |
| Post-dialysis | Full-range fit | $0.0007 \pm 0.000103$ | $8.69 \times 10^{-13}$ | 502 |
| | Restricted-range fit | $0.0008 \pm 0.000006$ | $9.75 \times 10^{-13}$ | 448 |
|  | Malvern NNLS | – | – | 615 (341–1106) |

### Supplementary Video

**Video S1**. Representative nanoparticle tracking analysis (NTA) video of rSP-B in 0.2% (w/v) DPC/SDS (9:1), showing the Brownian motion of individual particles during analysis. The corresponding particle size distribution is presented in Figure 2.

